# Mouse lemur, a new animal model to investigate cardiac pacemaker activity in primates

**DOI:** 10.1101/2021.10.25.465774

**Authors:** Mattia L. DiFrancesco, Romain Davaze, Eleonora Torre, Pietro Mesirca, Manon Marrot, Corinne Lautier, Pascaline Fontes, Joёl Cuoq, Anne Fernandez, Ned Lamb, Fabien Pifferi, Nadine Mestre-Francés, Matteo E. Mangoni, Angelo G. Torrente

## Abstract

**Background:** Gray mouse lemurs (*Microcebus murinus*) are the smallest and most ancestral primates known. Their size falls in between that of mice and rats, and their genetic proximity to humans makes them an emerging model to study age-related neurodegeneration. Since mouse lemurs replicate similar senescence processes of humans, they constitute a useful model for studying cardiovascular dysfunctions. However, their cardiac physiology is unknown. Thus, we investigated the cardiac pacemaker activity generated by the sinoatrial node (SAN) of mouse lemurs, presenting the first characterization of heart automaticity in nonhuman primates.

**Methods and Results:** We recorded cardiac automaticity in mouse lemurs and in their SAN tissues and pacemaker myocytes. Mouse lemurs have a heart rate (HR) in between those of mice and rats and a similar generation of the SAN electrical impulse. Their SAN myocytes express the main pacemaker currents at densities similar to mice: the hyperpolarization-activated current (*I_f_*) and the L-type (*I_ca,L_*) and T-type (*I_ca,T_*) calcium currents. Conversely, their ventricular depolarization resembles that of large mammals and despite the small size of mouse lemurs, the total number of heartbeats in their life corresponds to what can be attained by humans.

Using muscle-derived stem cells (MDSCs) from mouse lemurs, we also differentiated pacemaker-like (PML) cells showing spontaneous automaticity and expressing markers of native SAN myocytes (HCN4 and connexin-45).

**Conclusions:** Our characterization of heart automaticity in *Microcebus murinus* provides new opportunities for comparative cardio-physiology studies in primates and with humans and for testing molecules that could modulate age-related dysfunctions of heart rate.

## Introduction

The sinoatrial node (SAN) is the cardiac tissue that serves as the natural pacemaker of the heart.^1,2^ This tissue is situated at the venous inflow of the right atrium and is enriched in spindle-shaped cells that spontaneously depolarize to generate cardiac impulses.^2^ The pacemaker activity of SAN myocytes sets the intrinsic rate of heartbeats, which is constantly modulated by the autonomic nervous system to meet the physiological needs of the body.^3,4^

Aging is a major causes of dysfunction in SAN pacemaker activity.^5^ SAN dysfunctions and related brady-arrhythmias account for about half of the 500,000 electronic pacemaker implanted in Europe annually,^6^ costing $2 billion.^7^ Given these high costs and because of the increased incidence of SAN dysfunction in the aged population,^8^ complementary and alternative therapies to the implantation of electronic pacemakers are urgently required.^9^

This justifies the exploration of new species that are genetically closer to humans to better understand the mechanism of cardiac pacemaker activity. Grey mouse lemurs, *Microcebus murinus*, belong to the primate order and thus have common genetic origins with humans.^10,11^ Mouse lemurs are small and ancestral primates (Suppl. Fig 1A.) that typically live 4-5 years in the wild but can live up to 12 years in captivity, undergoing a period of prolonged aging like humans. During their life these small primates generate a total number of heartbeats similar to what can be attained by aged people (see discussion).

These lemurs are bred in captivity because, unlike rodents, they spontaneously develop age-related neurodegeneration similar to humans,^12–16^ which could also affect the autonomic nervous system. Thus, in comparison to rodents, mouse lemurs are a better animal model to closely replicate the nervous context that modulates heart activity as people age. Mouse lemurs also have some unique and ancestral features for primates, such as the ability to enter into a torpor state to conserve energy when the outside temperature drops.^17,18^

In summary, given the phylogenetic proximity to humans,^10,19^ similar neurodegenerative processes and equivalent number of heartbeat in a life, mouse lemurs are an ideal animal model for studying the effects of aging on heart pacemaker activity.

The mechanism of SAN pacemaker activity starts with the diastolic depolarization, a spontaneous phase that brings the membrane potential from the end of the repolarization phase to the threshold of the next action potential (AP).^20^ This mechanism has been extensively studied in rabbits and mice.^21–23^ Moreover, the introduction of transgenic mice has allowed researchers to gain relevant insights into the role of specific ion channels and proteins in the mechanism of pacemaker activity.^2,24^

Although several aspects of the generation of diastolic depolarization are still not fully understood, the evidence collected in rodents suggests that a functional association between plasma membrane ionic channels and intracellular Ca^2+^ dynamics generates the inward current required to depolarize SAN myocytes.^25^ Among various plasma membrane ionic channels, hyperpolarization-activated “funny” (f) channels encoding the f current (*I_f_*), L-type Ca_v_1.3 and T-type Ca_v_3.1 Ca^2+^ channels underlying *I_ca,L_* and *I_ca,T,_* are major mechanisms for generating the diastolic depolarization. As well, Ca^2+^ dynamics produced by the release of sarcoplasmic Ca^2+^ from the ryanodine receptors (RyRs) and Ca^2+^ efflux through the Na^+^/Ca^2+^ exchanger (NCX) generate another inward current (*I_NCX_*) that helps complete the diastolic depolarization of pacemaker cells.^25–27^

Despite the aforementioned evidence, the general mechanism of pacemaker activity in primates, and more specifically in our species, are still unclear. Indeed, it is difficult to study the living hearts of humans or other primates because of limited availability and strong ethical concerns. Therefore, only a reduced number of studies have been conducted to study the mechanisms of heart automaticity in the taxonomic order of primates^28–30^ and in humans.^31^ Thus, although the SAN pacemaker activity in mice closely resembles that in humans,^22,32,33^ mice only have 40% of the human genes required for viability, so the genetic distance must be taking in account when considering the mechanism of pacemaker activity.^10^

For that reason, here we investigated for the first-time heart automaticity in mouse lemurs. These primates have an average weight of 60–90 g and a body length of 10–12 cm (Suppl. Fig. 1A).^34^ Since the frequency of heart contractions in mammals is determined by their body size, in order to sustain physiological and metabolic needs,^35,36^ we compared our results on mouse lemurs to previous findings on mammals of similar sizes, such as rats and mice. We also compared the cardiac activity in mouse lemurs and humans when possible and relevant.

For this purpose, we characterized the cardiac pacemaker activity of mouse lemurs in three different settings: (i) freely moving and anesthetized animals, (ii) isolated hearts and SAN tissue preparations, and (iii) single SAN pacemaker myocytes. In addition, we used biopsies of mouse lemur skeletal muscles to isolate muscle-derived stem cells (MDSCs). Then, we differentiated *in vitro* MDSCs into automatic cells with a pacemaker-like (PML) phenotype similar to native pacemaker myocytes of the SAN, as we previously reported in rodents.^37,38^

In conclusion, our general characterization of cardiac automaticity in mouse lemurs paves the way for future studies of age-related SAN dysfunctions in primates, making it easier to translate basic research findings into clinical applications against cardiovascular diseases.

## Results

### 1. Analysis of the heart rate (HR) of freely moving and anesthetized mouse lemurs

We first used subcutaneous telemetry transmitters to capture electrocardiograms (ECGs) in mouse lemurs aged 1–5 years. This technique allowed us to discern P waves, QRS complex, and T waves, which correspond to atrial contraction, ventricular depolarization, and ventricular repolarization, respectively^39^ (Fig. 1A). After analyzing ECGs recorded in one mouse lemur for 22 h, the main heart rate (HR) range was found to be between 250 and 550 bpm (Fig. 1B), while the average HR was of 427±43 bpm (n=4; Fig. 2B). This range coincides with the HR reported for rodents of similar size such as mice^40^ and rats,^41,42^ respectively, and is consistent with the body size of mouse lemurs. In addition, a graphical representation of the HR over time in two mouse lemurs revealed HRs as high as 520 bpm and as low as 160 bpm (Fig. 1C). In particular, the gradual slowing of the HR to about 200 bpm appeared to be a recurring pattern, often followed by a sudden increase in HR (Fig. 1C and Suppl. Fig. 1). HR variability (HRV) analysis revealed that the periods of HR slowdown had a slow dynamic that differed from short-term HRV generated by parasympathetic regulation of SAN pacemaker activity.^4^ Indeed, such an HR slowdown influenced more the long-term HRV indexes like the coefficient of variability of the RR interval between successive heartbeats, rather than short-term HRV indexes (root mean square of successive differences [RMSSDs] between heartbeat intervals; Fig 1D). We obtained similar conclusions from Poincaré analyses of the short-term and long-term HRV indexes (SD1 and SD2 respectively; Suppl. Fig. 1).

**Figure 1:**
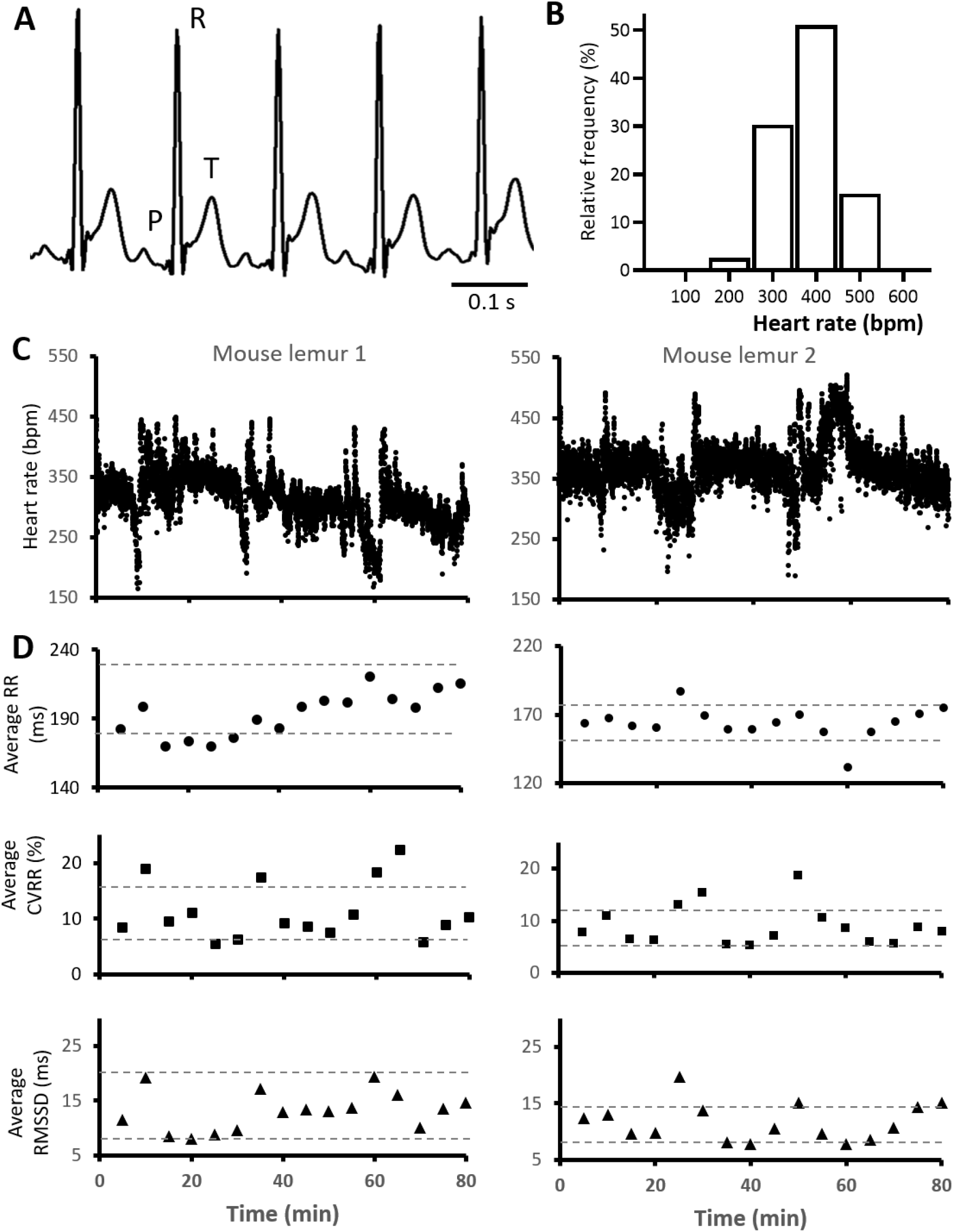
*In vivo* recording of heart rate in mouse lemurs. **A:**Sample traces of ECGs in freely moving mouse lemurs. Note P, R and T waves. **B:**Range of heart rates (HR) measured during 22 hours of ECG recording in one mouse lemur. Bars represent the percentage of binned heart rate values. Binning width is 100 bpm. **C:**Example of 80-min recordings in two mouse lemurs showing HR changes over time. Dots represent consecutive R-R intervals. Note cyclical HR decrease to very low frequencies. **D:**Plots obtained by averaging every 5 min the interval between successive heartbeats (RR interval), its coefficient of variability (CVRR) and the root mean square of successive differences between heartbeats (RMSSD), for the recordings shown in C. Dashed gray lines indicate the threshold of mean ± SD for each of the parameters reported in this panel.

**Figure 2:**
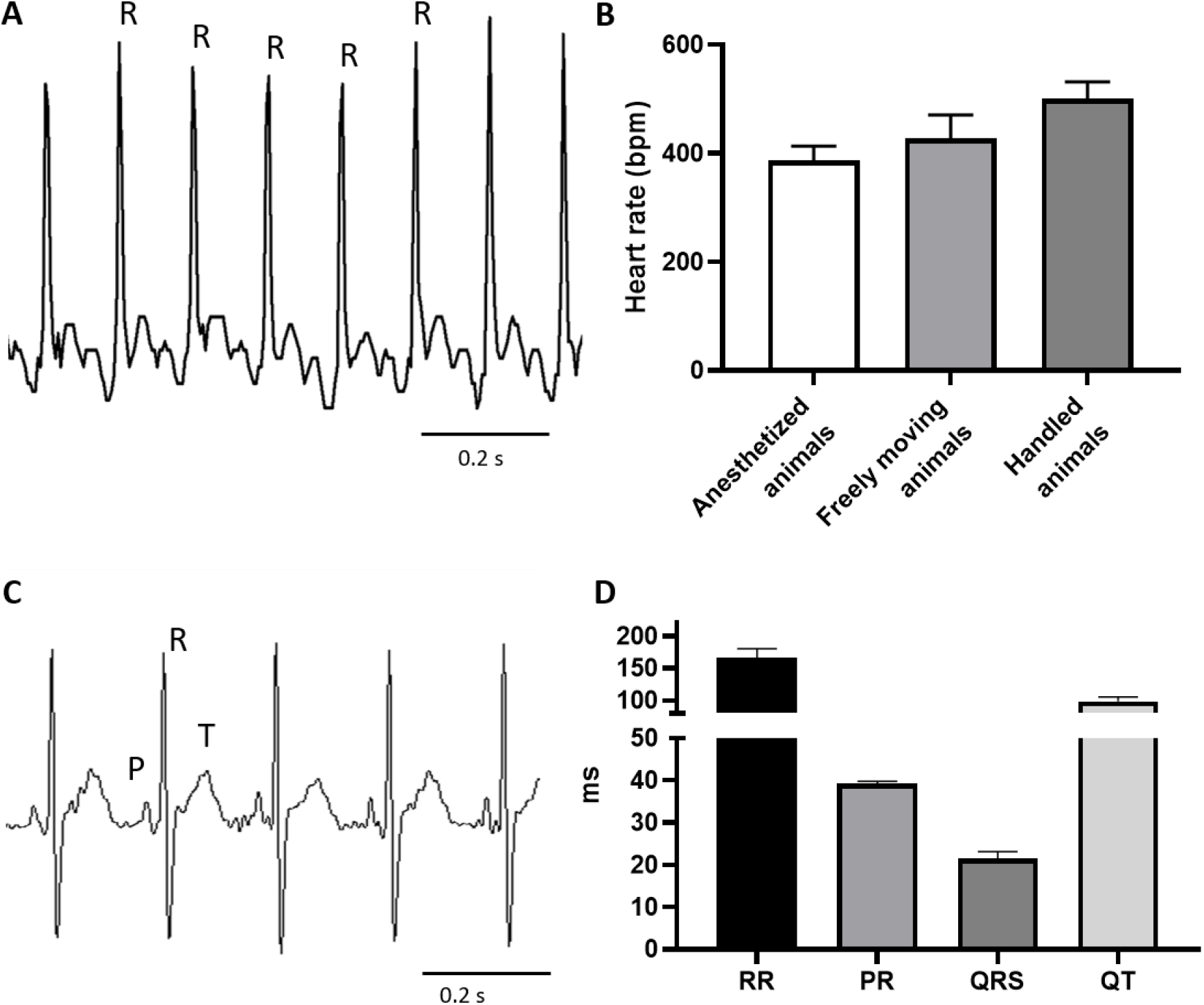
Cardiac activity of mouse lemurs during handling or anesthesia. **A:**Sample of surface ECG in mouse lemurs recorded with an external device just after handling of the animals. **B:**HR comparison in anaesthetized, freely moving and handled mouse lemurs (n=5, n=4, n=3, respectively). **C:**Electrocardiograms from anaesthetized mouse lemurs. Note P, R and T waves. **D:**interval between successive heartbeats (RR), interval between atrial and ventricular contraction (PR), length of the ventricular depolarization (QRS) and interval between ventricular depolarization and repolarization (QT) characterizing the ECG waveform of anaesthetized mouse lemurs (n=5).

In mouse lemurs’ ECGs we also observed visible T waves separated from the QRS complexes by short isoelectric ST segments (Fig. 1 A). These T waves differ significantly from those of small rodents^39^ and are more similar to those recorded in large mammals^43^ and humans.^41,44^

Next, we used an external ECG device to obtain shorter recordings of the HR of mouse lemurs. In these experiments, the sympathetic tone was likely increased by gentle handling of the animals to put the external device in place. Indeed, this method revealed an average HR higher than animals equipped with subcutaneous ECG transmitters, consistently with the expected response to the stimulation of the sympathetic pathway (Figs. 2A and B).

We also recorded ECGs of the anesthetized animals, which allowed us to measure the cardiac activity under reduced autonomic input (Figs. 2C and D). The average HR determined under these conditions was 372 ± 27 bpm (n = 5; Fig. 2B), which places the HR of mouse lemurs between those of mice and rats according to mouse lemurs body size.^40,41^ We identified T waves separated from the QRS complex also in the anesthetized mouse lemurs, which is consistent with recordings obtained from freely moving animals.

### 2. Analysis of the intrinsic HR of the isolated hearts and optical mapping of the intrinsic SAN impulse in the intact atrial preparations

To comply with the “three Rs”of animal welfare (Replacement, Reduction and Refinement), we obtained results on the isolated cardiac tissues of mouse lemurs only from animals aged 6–12 years that had to be euthanized because of natural ageing or other factors interfering with normal life. Thanks to this approach we were able to record the ECGs of isolated hearts of mouse lemurs. Their hearts were obtained after deep anesthesia of the animals and complete loss of hind-limb reflex or any other sensation. After removal, the hearts were placed in a Langendorff perfusion system that maintained an average HR of 232 ± 80 bpm (Fig. 3 A). Such a HR was similar to what we recorded by optical mapping in intact SAN tissues (Fig. 3). To obtain these tissue preparations, which included the SAN and the right and left atria, we removed the ventricular chambers in hearts isolated from other deep anesthetized animals. The SAN preparations of mouse lemurs were about twice the size of those observed in mice of comparable age (24 months old; Fig. 3B). Moreover, while the left atria of aged mice were slightly smaller than the right atria, the left atria of aged mouse lemurs were significantly larger than the right atria (Suppl. Fig. 2), suggesting an age-related remodeling of the heart’s chambers in mouse lemurs.

**Figure 3:**
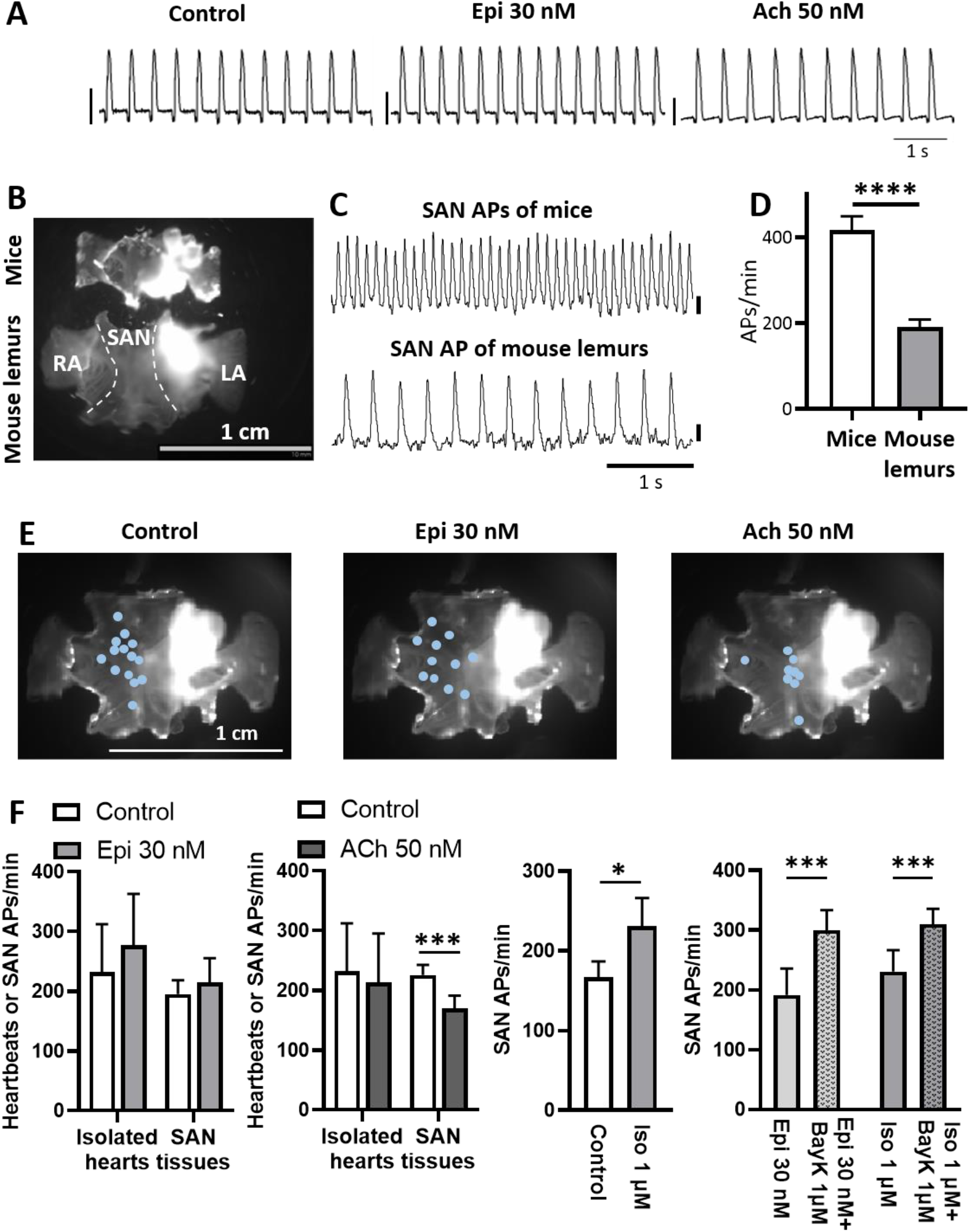
Pacemaker activity of isolated hearts and SAN preparations. **A:** Sample traces of ECGs recorded in isolated hearts from mouse lemur, maintained under constant perfusion with a Langendorff system. Traces have been recorded under control conditions (perfusion with Tyrode’s solution), perfusion of epinephrine (Epi; 30 nM) or acetylcholine (Ach; 50 nM). Scale bars correspond to 1 mV. **B:** Examples of SAN preparations of mice and mouse lemurs, including the SAN, the right and left atria (RA and LA, respectively). **C:** Example traces of optical action potentials (APs) recorded in the SAN of mice and mouse lemurs. APs amplitude is expressed in arbitrary units of fluorescence (a.u.), scale bars correspond to 1 a.u. **D:** Rate of APs recorded in mice and mouse lemurs under control condition (n mice = 13, n mouse lemurs = 17). **E:** Dots indicating leading pacemaker sites in SAN preparations of mouse lemur. **F:** Change in heart rate (heartbeats or APs/min), before and after Ach, Epi or isoprenaline (Iso; 1 μM) superfusion, or during Epi and Iso superfusion and after addition of BayK8644 (BayK; 1μM), in Langendorff perfused hearts (n=3) or atrial preparation of mouse lemurs (n=9 in Epi, n=11 in ACh, n=10 in Iso, n=5 in Epi and Epi + BayK and n=10 in Iso and Iso + BayK). *p<0.05, ***p<0.001 and ****p<0.0001 by unpaired T-test, paired T-test, two-way Anova and two-way Anova with Sidak’s multi comparison test.

We used optical mapping of membrane voltage to record the rate of spontaneous AP generation in the SAN region of both mouse lemurs and mice, as well as the localization of the origin of pacemaker depolarization and the conduction of the electrical impulse from the SAN to the atria (Figs. 3 and Suppl. Video 1). We discovered that the average pacemaker frequency in the mouse lemurs SAN was about half of what we recorded in mice of comparable age (24 months old; Figs. 3C and D), but was faster than the AP rate observed in aged rats.^42^

We also tested the chronotropic response of isolated hearts and SAN tissues to natural agonists of sympathetic and parasympathetic nervous systems. When we superfused the β-adrenergic agonist epinephrine (30 nM), we detected only a tendency toward positive chronotropic modulation. This outcome could be related to the advanced age of the animals used, which could partially impair the β-adrenergic response to epinephrine (Figs. 3A and F). Conversely, the vagal effector, acetylcholine (50 nM), induced a significant rate reduction in the same preparations (Figs. 3A and F). For both the adrenergic and vagal agonists, the modulation of the SAN rate in the tissue preparations was associated with a shift in the origin of pacemaker activity toward the frontal or caudal portion of the SAN (Fig. 3E), respectively, similar to our previous observations in mice.^45^

We further investigated the β-adrenergic response using saturating doses^46^ of the artificial sympathetic agonist, isoprenaline (1 μM), on SAN tissues analyzed by optical mapping. In comparison to epinephrine (30 nM), this drug amplified the response of the SAN to the β-adrenergic stimulation, causing a significant increase in the generation of spontaneous APs (Fig. 3F). In addition, direct activation of the L-type Ca^2+^ channels with the dihydropyridine agonist Bay K 8644 (1 μM), induced a much stronger increase of AP rate in SAN preparations previously exposed to β-adrenergic stimulation with epinephrine (30 nM) or isoprenaline (1 μM; Fig. 3F). The stronger AP rate increase under Bay K 8644 (1 μM) indicates that direct activation of L-type Ca^2+^ channels in aged mouse lemurs was more effective in stimulating the chronotropic response of pacemaker activity in comparison with β-adrenergic stimulation.

### 3. Pacemaker activity and ionic currents in native pacemaker myocytes isolated from the SAN of mouse lemurs

Individual pacemaker myocytes were isolated from the SAN tissue of mouse lemurs to characterize the SAN pacemaker activity at the cellular level. After enzymatic digestion of the mouse lemur SAN, we observed single spindle-shaped pacemaker myocytes with self-contractile properties (data not shown), which are similar to SAN pacemaker myocytes in mice,^2^ rats,^47^ and humans^31^ (Fig. 4A).

**Fig 4:**
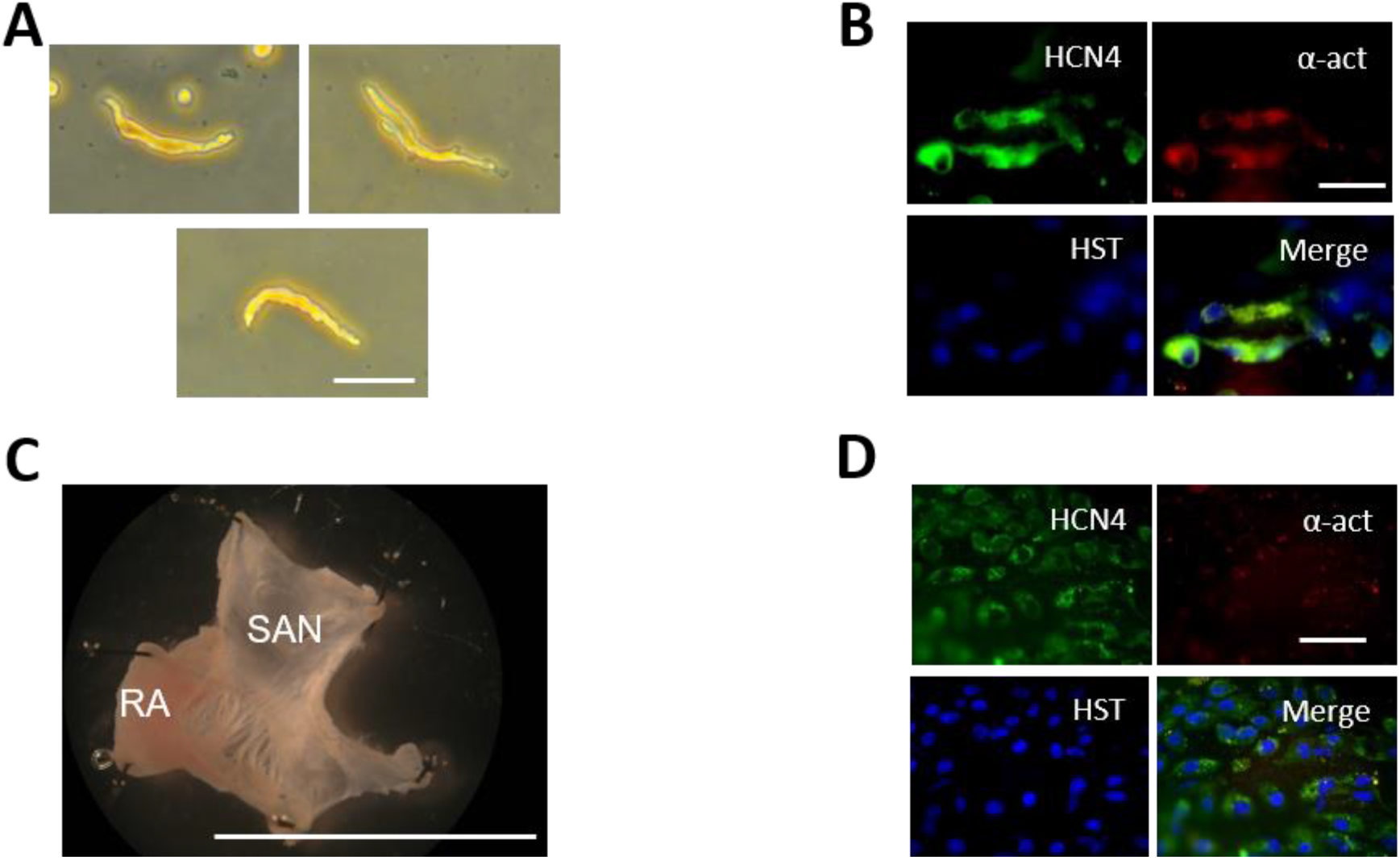
Immuno-detection of cardiac-pacemaker markers from mouse lemur SAN tissue and isolated myocytes. **A:**Freshly isolated pacemaker myocytes obtained from mouse lemur SAN tissues (scale bar = 50 μm). **B**: Labelling of HCN4, α-actinin and nuclei (Hoechst33358; HST), together with merged picture in SAN pacemaker myocytes (scale bar = 30 μm). **C**) Dissection of the atrial preparation used to isolate SAN myocytes, showing the SAN and the right atrium (scale bar = 10 mm). **D**) Labelling of HCN4, α-actinin, nuclei (HST) and merged picture of pacemaker myocytes within the intact SAN (scale bar = 40 μm).

The immunostaining of single pacemaker myocytes and intact mouse lemur SAN tissues revealed the expression of HCN4 proteins, which encode f-channels and are one of the main markers of pacemaker myocytes (Figs. 4B, C, and D). According to the molecular presence of f-channels, in pacemaker myocytes isolated from the mouse lemur SAN we recorded the funny current (*I_f_*) at a density similar to that of mice,^48^ i.e., 2.5-fold higher than human SAN myocytes.^31^

In mouse lemur SAN myocytes we also recorded the L-type and T-type Ca^2+^ currents, *I_ca,L_* and *I_ca,T_*, which, together with *I_f_*, are among the main currents involved in the mechanism of pacemaker activity (Figs. 5A and B). Mouse lemur SAN myocytes had a lower total Ca^2+^ current (*I_ca,Tot_* = *I_ca,L_* + *I_ca,T_*) than mouse myocytes, possibly because of the lower *I_ca,T_*. However, *I_ca,L_* density and its activation threshold were similar to that in the SAN myocytes of mice.^49^ Conversely, *I_ca,L_* of mouse lemurs SAN myocytes peaked at 0 mV, which is more positive than the peak current in mice (−10 mV)^50^ but similar to that reported in rat.^47^ In addition, in mouse lemurs SAN myocytes we recorded the inward-rectifier K^+^current *I_K1_* (Figs. 5A and B) at a density similar to what has been recorded in rats.^47^

**Figure 5:**
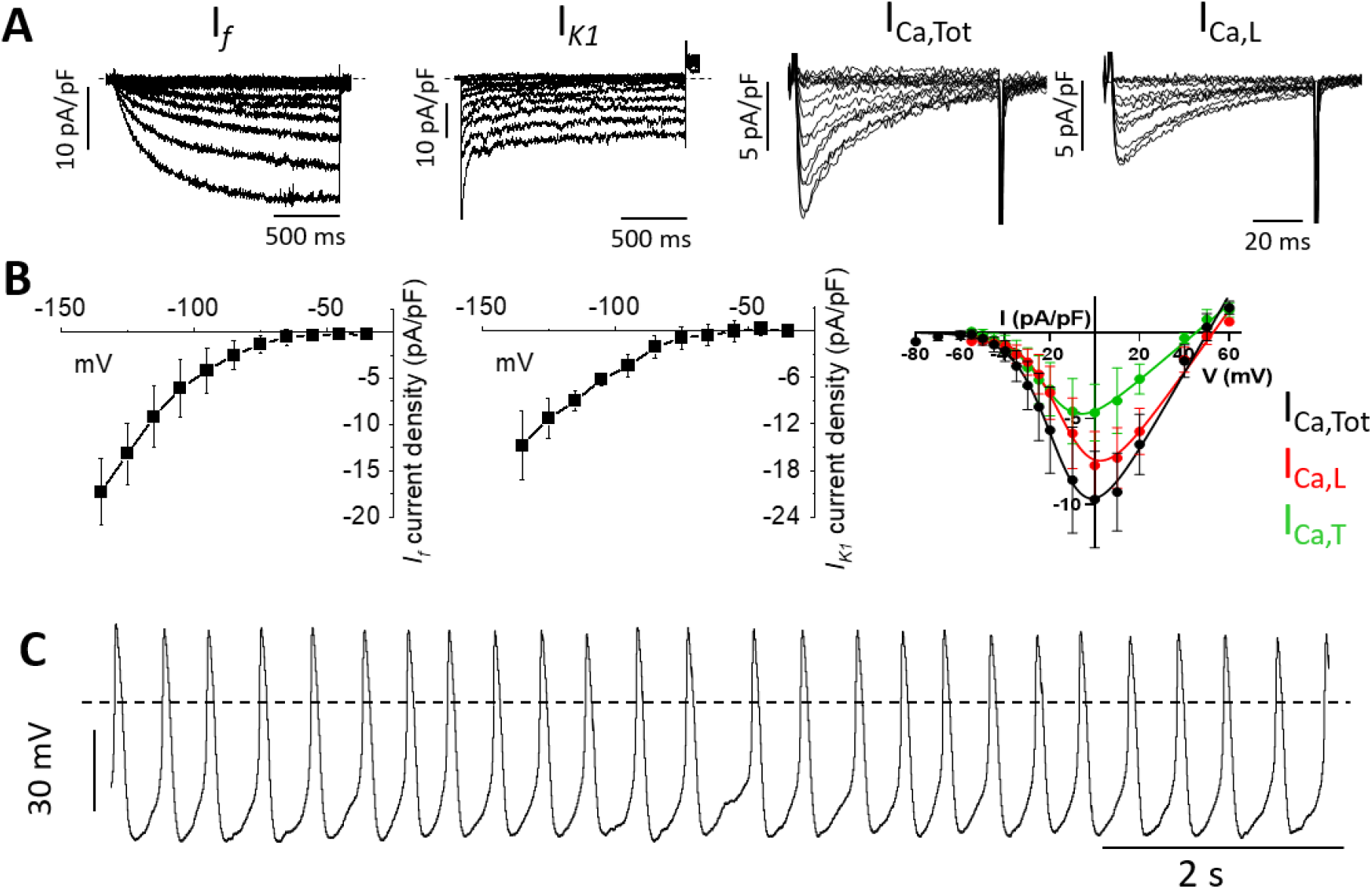
Recordings of ionic currents in SAN myocytes of mouse lemur. **A:**Sample traces of *I_f_, I_K1_*, (n = 4 and 3), total *I_Ca_ (I_ca,Tot_*) and *?ca,L* (n = 4). **B:**Current-to-voltage relationships of densities of ionic currents described in panel (A) for *I_f_*, *I_K1_, I_ca,Tot_, I_ca,L_* and *I_ca,T_*.**C:**Action potentials (APs) recorded in SAN pacemaker myocytes from *Microcebus murinus* under perfusion of Tyrode’s solution.

Current clamp recordings of the pacemaker activity in mouse lemur SAN myocytes under Tyrode’s solution revealed spontaneous APs with an average beating rate of 148 ± 22 APs/min (n = 7; Fig. 5C).

### 4. Analysis of automaticity in PML cells differentiated from MDSCs

Although small primates such as mouse lemurs can be kept in captivity for fundamental research and species conservation purposes, their availability remains limited in comparison to that of classical rodent models such as mice or rats. Such a limited availability reduces the possibility to study pacemaker activity in mouse lemurs at the tissue or cellular level. Conversely, *in vitro* preparations of stem cells are being increasingly used for studying cellular physiology and ion channel function.^51^

We previously demonstrated that cells with similar characteristics to native SAN pacemaker myocytes can be differentiated from adult skeletal MDSCs of mice.^37,35^ Therefore, we developed an *in vitro* model of cellular automaticity from mouse lemurs, by using skeletal muscle biopsies that we differentiated in cells that recapitulate many of the characteristics of native SAN pacemaker myocytes. For that purpose, we used the pre-plating technique to isolate the undifferentiated MDSCs, which were then cultured for 66–74 days, to ensure a full differentiation toward the sustained beating phenotype of PML cells.^38^

The mouse lemur PML cells did not express myogenin, indicating that they do not belong to the myogenic lineage typical of skeletal muscle cells (Fig. 6A). Similar to native pacemaker myocytes of the murine SAN, mouse lemur PML cells co-expressed connexin45 (C×45) with alpha-actinin and caveolin-3. The C×45 positive cells also co-expressed L-type Ca_v_1.3 Ca^2+^ channel proteins (Fig. 6A). In addition, we observed co-expression of the f-channels (HCN4) with cardiac troponin C (Fig. 6A). These data showed that PML cells differentiated from mouse lemurs MDSCs exhibit typical markers of native SAN pacemaker myocytes.^2^

**Figure 6:**
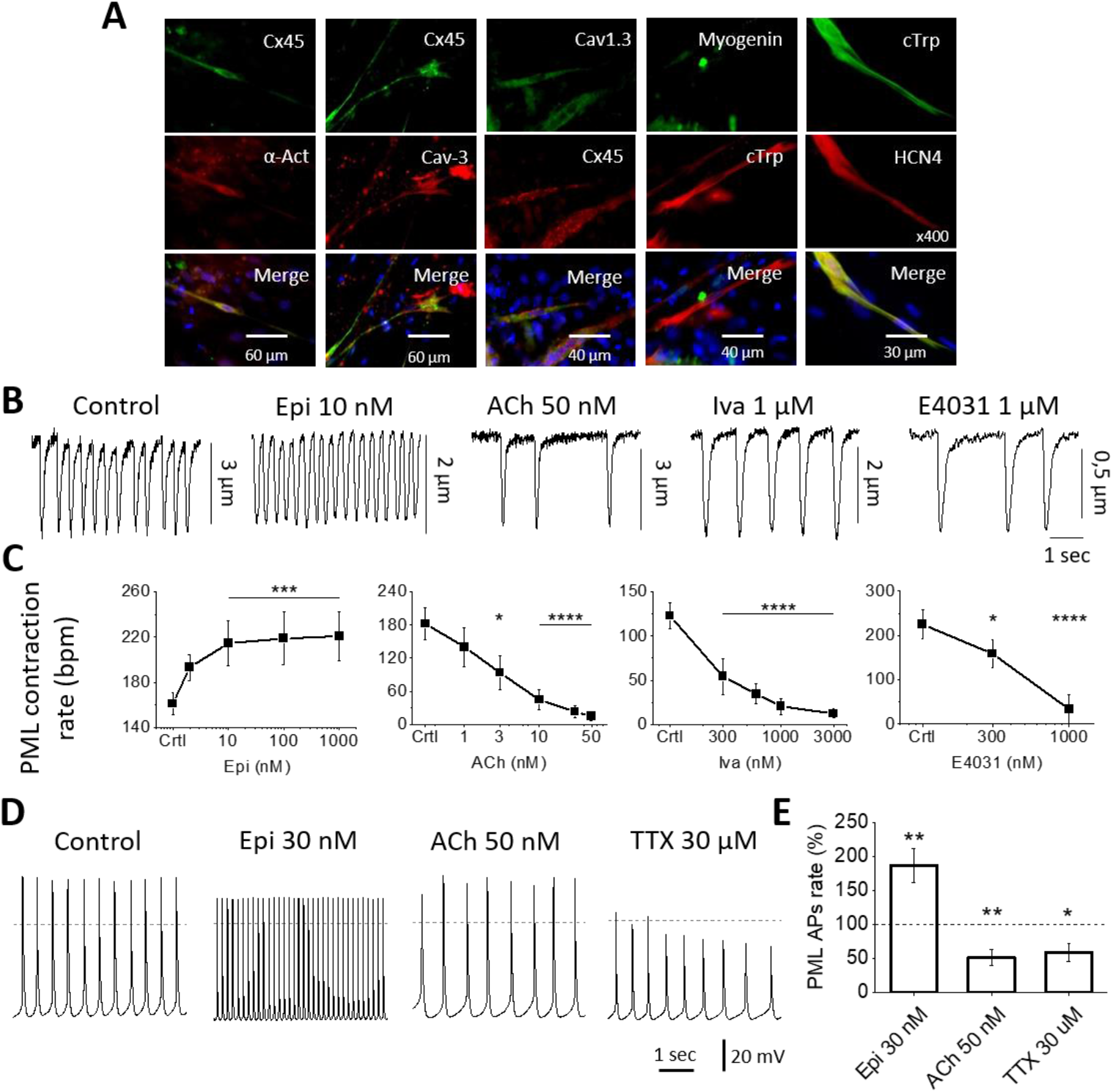
Automatic profile of PML cells derived from mouse lemur skeletal muscles. **A:**Immunostaining of PML cells with the SAN marker HCN4, Cx45 and Cav1.3 plus sarcomeric alpha-actinin (α-Act), Caveolin 3 (Cav-3), myogenin and cardiac troponin I (cTrp). **B:**Example traces of cell contractions recorded in PML cells under control and after modulation with epinephrine (Epi), acetylcholine (Ach), ivabradine and E4031. **C:**Dose-response of contraction rate from PML cells under the condition in B (n = 7, 11, 9, 6 respectively). **D:**Example traces of APs recorded in PML cells under control and after modulation with epinephrine, acetylcholine and TTX. **E:**% change of spontaneous APs frequency under the conditions in D. Dashed line indicates AP rate in control conditions (n = 9, 8, 6 respectively). Paired Wilcoxon’s t-tests were applied between normalized internal control and values after drug perfusion. *p<0.05, **p<0.01, ***p<0.001 and ****p<0.0001 by Two-way ANOVA.

Similar to native pacemaker SAN myocytes, the PML cells contracted spontaneously at a regular rhythm that could be detected by an edge-detection system. Thus, our PML cells were in a physiologically favorable condition for dose-response analysis of modulator compounds. To this end, we used spontaneous contraction of the PML cells to assess the effects of increasing doses of epinephrine, acetylcholine, ivabradine, and E4031, an inhibitor of the fast component of the delayed rectifier K^+^ current (*I_Kr_*). The PML cells had a spontaneous contractility rate of 171 ± 13 contractions/min (n = 33; Fig. 6B) under control conditions, which is equivalent to the AP firing measured in native mouse lemur pacemaker myocytes (Fig. 5C). The β-adrenergic agonists epinephrine accelerated the spontaneous activity of the PML cells in a dose-dependent manner. Conversely, the activation of the muscarinic receptors by Ach, slowed automaticity accordingly to the doses used (Figs. 6B and C). The *I_f_* blocker (ivabradine) and the *I_kr_* inhibitor (E-4031) decreased the rate of spontaneous contractions of the PML cells, indicating the functional expression of HCN4 channels and of the ERG K^+^ channels responsible of *I_kr_*, respectively (Figs. 6B and C).

Current-clamp recordings of the PML cells revealed spontaneous AP firing, with a rate of 161 ± 14 APs/min (n = 12; Fig. 6D), which is similar to what we recorded in the native cardiac pacemaker myocytes of mouse lemurs (Fig. 5C). The modulation of spontaneous APs by β-adrenergic and muscarinic receptors was evaluated by superfusing epinephrine and ACh. The firing rate of the PML cells increased by 87 ± 25% under β-adrenergic stimulation with epinephrine (30 nM), while it decreased by 49 ± 11% under Ach superfusion (50 nM; Fig. 6D and E). In addition, we tested the functional presence of the fast Na^+^ current (*I_Na_*) by superfusing tetrodotoxin (TTX). TTX (30 μM) significantly decreased the spontaneous beating rate of the PML cells by 41 ± 13% (Figs. 6D and E).

## Discussion

*Microcebus murinus* (Suppl. Fig 1A.) is an arboreal and nocturnal primate that is among the smallest and most primitive of its order.^52^ This lemur is emerging as a new model for studying neurodegenerative diseases related to aging, because it better recapitulates the neurophysiology of humans.^12–16^ Given the importance of the heart-brain crosstalk in HR modulation^53^ and the phylogenetic proximity of *Microcebus murinus* to *Homo sapiens*,^10^ this lemur could also help the research in cardiovascular physiology.^52^ In addition, mouse lemurs can be bred in captivity because of their small size and easy reproduction, allowing to maintain a viable population that could help preserve these lemurs for future reintroduction into the wild, to fight against the current threat of habitat loss.

Abnormal regulation of heart pacemaker activity by the autonomic nervous system has been related to aging.^23,54^ Similar to humans, mouse lemurs spontaneously generate neurodegenerative diseases as they age, a feature that is not found in mice.^13^ This degeneration possibly affects the ability of the autonomic nervous system to regulate cardiovascular functions. Thus, mouse lemurs can satisfactorily recapitulate a neural environment similar to that in humans, allowing for comparative studies of cardiac automaticity and age-related dysfunctions. In addition, given the average HR of about 430 bpm that we found in mouse lemurs and a lifespan of 12 years, their heart beats a total of 2.7 billion of time in a life which is very close to the 3 billion of heartbeats generated by a person of 80 years with an average HR of 72 bpm. Conversely, even considering 3 years of maximal age for mice and an average HR of 490 bpm,^40^ their heart beats only 0.8 billion times in a life. Likewise, considering an HR of 350 bpm in rats^42^ and a maximum lifespan of 4 years, their heart beats only 0.7 billion times in a life. Thus, humans and mouse lemurs generate about three times more heartbeats in a life than mice or rats.

However, despite the advantages of using mouse lemurs for comparative studies of cardiac physiology, no research on the heart pacemaker activity of these lemurs has been published to date. Thus, to study the cardiac physiology of mouse lemurs, we used a range of techniques to characterize the rate and pacemaker activity of the heart and compare them to previous data from similar-sized rodents (mice and rats) or humans.

Our experiments revealed that the HR of mouse lemurs falls in-between the rate previously reported in mice and rats.^40,42^ These data are consistent with the small size of this primate. Conversely, despite the small size of mouse lemurs, the total number of heartbeats in their life (2.7 billion) is much closer to humans (3 billion) than to mice (0.8 billion) or rats (0.7 billion). Moreover, the ECGs of mouse lemurs displayed T waves that were visibly separated from the QRS complex, suggesting repolarization dynamics more similar to that of large mammals^43^ and humans,^41,44^ which further distinguish cardiac features in mouse lemurs from small rodents.

In mouse lemurs, we also observed recurrent reductions in the heartbeat to 200 bpm or less, which were frequently followed by a sudden increase in HR. The HR oscillates at a fast time scale of less than a second when the parasympathetic branch of the autonomic nervous system is activated/desactivated.^4^ The long-time scale of HR decrease in the mouse lemurs suggests that this is a different phenomenon. We do not have an immediate explanation for the HRV dynamics, but the long time scale suggests that the decrease in HR could be a physiological phenomenon involving more complex causes than classical parasympathetic activation.^4^ We hypothesize that this slow HR decrease could be related to the mechanism that allows mouse lemurs to enter into a torpor state to conserve energy when the outside temperature drops.^17,18^

Our analysis of the atrial preparations of mouse lemurs revealed that the size of the supraventricular region was about twice that of mice. Conversely the intrinsic frequency of AP generation in the SAN of mouse lemurs was about half of what we recorded in equivalent preparations of mice, in accordance with the difference in size. Moreover, we discovered that the left atria of mouse lemurs were larger than the right atria. This difference could be due to the old age of the mouse lemurs used for these experiments, and it could also be a point of similarity with human cardiac physiology. Indeed, it is well known that as people age, their left atria enlarge to compensate for a lack of left ventricle contraction, helping ensure adequate blood circulation.^55^ The same compensatory remodeling of the heart’s chambers might take place in aged mouse lemurs.

Studying single pacemaker myocytes isolated from the mouse lemur SAN we demonstrated the expression of HCN4 channels as well as the conduction of *I_f_*. In addition, we observed an early activation of *I_ca,L_* at negative voltages, similar to what we discovered in murine SAN myocytes that express Ca_v_1.3 channels.^49^ The conduction of early activated *I_ca,L_* in the SAN of mouse lemurs, and the positive chronotropic response to the *I_ca,L_* agonist Bay K 8644, suggest that Ca_v_1.3 channels play an important role in the pacemaker mechanism of primates. Such an hypothesis is consistent with the recent discovery of inherited channelopathies of *Cacna1D* genes, responsible of Ca_v_1.3 expression, which cause SAN dysfunction in humans^56^ and shed light on the important role played by Ca_v_1.3 channels in the generation of pacemaker activity^21,50^.

Similar to previous reports on SAN myocytes from mice^57^ and rats,^47^ we also discovered Ba^2+^-sensitive *I_K1_* in the mouse lemurs at densities comparable to those observed in rat SAN myocytes.^47^ The *I_K1_* is almost absent in the SAN myocytes of relatively large mammals, but it is expressed in the working myocardium and Purkinje fibers, where it is responsible for the negative diastolic potential compared to SAN myocytes.^2^ Thus, the presence of this current in mouse lemurs could indicate a specificity of small mammals.

We discovered that the natural β-adrenergic agonists epinephrine at a dose of 30 nM only induced slight positive chronotropic effects on SAN preparations, or intact hearts of aged mouse lemurs. Conversely, saturating β-adrenergic stimulation with the artificial agonist isoprenaline significantly increases the frequency of pacemaker activity in the SAN. In addition, further stimulation with the *I_ca,L_* agonist (Bay K 8644) causes a much stronger increase of the SAN AP firing in comparison with previous stimulation with epinephrine (30 nM) or isoprenaline (1 μM). Such results suggest that the sympathetic pathways could be impaired by the old age of the mouse lemurs, needing a saturating dose of β-adrenergic agonists to be significantly activated. In addition, the much stronger chronotropic response to direct stimulation of pacemaker activity via L-type channels activation^49,58^ suggests that the final targets of the β-adrenergic stimulation of the SAN are still functional, regardless of the age of mouse lemurs. Concerning the muscarinic modulation, we found that acetylcholine superfusion significantly decreased the pacemaker rate in the tissue preparation tested.

Finally, we obtained MDSCs from the skeletal muscle biopsies of mouse lemurs and we differentiated them into PML cells using a technique that we had previously applied on mice.^37,38^ Our experiments showed that PML cells share similar features with native SAN pacemaker myocytes isolated from the mouse lemur SAN. In our recordings, PML cells contracted spontaneously and generated spontaneous APs at a rate comparable to that of native mouse lemur pacemaker myocytes. The PML cells expressed typical markers of native pacemaker myocytes, namely HCN4, Ca_v_1.3, and Cx 45. In addition, PML cells responded to positive and negative β-adrenergic and muscarinic chronotropic stimulations, respectively, as well as to *I_f_, I_Kr_* and *I_Na_* inhibitors.

Further studies will be required to better understand the degree of similarity between PML cells and native SAN myocytes.^38^ Nevertheless, our results suggest that PML cells could represent a valid model for a preliminary investigation of cardiac pacemaker activity in primates to evaluate molecules of pharmacologic interest.

In conclusion, we provide the first characterization of cardiac pacemaker activity in mouse lemurs and in their primate order. This characterization will help further research in cardiovascular dysfunctions associated with aging in a neurodegenerative context that is specific to primates and that effectively replicate that of humans. Thus, mouse lemurs will make it easy to apply basic findings on pacemaker activity to human physiology.

Moreover, this first characterization will aid future research into the mechanism underlying the torpor state of mouse lemurs. Although their ability to enter the torpor state may appear anecdotal at first, it could be very useful in inducing artificial comas, preventing organ failure, and slowing down human metabolism during long-term space flight.^18^

## Methods

### 1. Care and use of animals

This investigation conformed to the Guide for the Care and Use of Laboratory Animals published by the US national Institute of Health (NIH Publication No. 85–23, revised 1996) and European directives (2010/63/EU). Male and female *Microcebus murinus*, of 1 to 11 years of age, were used. Specifically, animals from 1 to 5 years were used for telemetry recording with minimally invasive or external electrocardiogram devices that allowed reintegration of the animals to their colony after the experimentation. Conversely, animals from 6 to 11 years, that after natural ageing had to be euthanized for factors hampering normal life, like weight loss, were used for all the other experiments. This approach reduces the need of animals in the respect of the “three R’s” of animal welfare: replacement, reduction, and refinement.

Mouse lemurs used for the implantation of electrocardiogram transmitters were hosted at the mouse lemur colony of Brunoy (MECADEV, MNHN/CNRS, IBISA Platform, agreement F 91.114.1, DDPP, Essonne, France) under the approval of the Cuvier Ethical Committee (Committee number 68 of the “Comité National de Réflexion Éthique sur l’Expérimentation Animale”), under authorization number 68-018. In the respect of the principle of “three R’s” of animal welfare, the use of these data, collected during another project for other purposes, allowed reduction of animal use since we did not have to implant new animals for the present project. All the other mouse lemurs came from the colony housed at RAM-CECEMA (license approval 34-05-026-FS, DDPP, Hérault, France). Their use was conducted under the approval of the CEEA-LR Ethical Committee (Committee number 36). Animals were euthanized at the end of their lives and all organs were collected immediately for biomedical research.

### 2. Housing conditions

All animals were housed in cages equipped with wood branches for climbing activities, as well as wooden sleeping boxes mimicking the natural sleeping sites of mouse lemurs. The temperature and the humidity of the rooms were maintained at 25–27 °C and at 55–65%, respectively. In captivity, the artificial daily light-dark cycles within the housing rooms are changed to mimic season alternation, with periods of 6 months of summer-like long days (14 h of light and 10 h of darkness, denoted 14:10) and 6 months of winter-like short days (10 h of light and 14 h of darkness, denoted 10:14). Animals were fed ad libitum with fresh fruit, worms and a homemade mixture as in Hozer et al.^59^

### 3. Electrocardiogram recordings

One-lead electrocardiograms were recorded from freely moving and anaesthetized mouse lemurs. For electrocardiogram recordings in freely moving animals implanted with a transmitter, surgery was conducted in sterile conditions, under veterinary supervision. After administration of diazepam (Valium, 1 mg/100 *g*, i.m.) and buprenorphine (0.005 mg/100 *g*, i.m.), anesthesia was induced and maintained by 1–3% isoflurane inhalation. Body temperature was maintained with a heating pad, and the animal’s eyes were protected with ocular gel (Ocry-gel; Laboratoire TVM, Lempdes, France). A small transmitter (PhysioTel F20-EET, 3.9 *g*, 1.9 cc; Data Sciences International TM, DSI, St. Paul, United States) connected with 1 pair of electrode wires (silicon elastomer insulated stainless-steel wires, diameter: 0.3 mm) was inserted inside the peritoneal cavity of the animal. Electrode wires were led subcutaneously from the abdomen to the thorax muscle and were sutured using non-absorbable polyamide monofilament suture thread, similarly to the procedure described in mice.^40^ After surgery, nociception was minimized by subcutaneous injection of analgesic and anti-inflammatory drugs (meloxicam, 0.2 mg/100 *g*). Electrocardiogram signals were recorded using a telemetry receiver and an analog-to-digital conversion data acquisition system (Data Sciences International TM, DSI, St. Paul, United States). HR and HRV data were analyzed with the dedicated software Labchart 8 (ADInstruments, Dunedin, New Zealand) and ecgAUTO v3.3.5.12 (emka TECHNOLOGIES, Paris, France).

For external electrocardiograms, we used gel electrodes (Youpin Hipee disposable electrocardiogram electrode patch) and the Xiaomi Youpin HiPee Intelligent Dynamic Holter produced by Zhejiang Helowin Medical Technology Co., Ltd (Hangzhou, China), to obtain one lead recording of HR. Soft surgical tape helps to keep the Holter on the back of mouse lemurs. Before that, the animals were shaved in the contact points with the gel electrodes, to facilitate the conduction of the signal through the skin. Electrocardiogram recordings were obtained through dedicated application software provided by the Helowin company and analyzed with the dedicated software Labchart 8 (ADInstruments, Dunedin, New Zealand). 5 min of electrocardiogram recordings obtained in the first 15 min after the application of the Holter were used to obtain HR just after handling of the animals.

For electrocardiogram recordings under anesthesia, we constantly exposed mouse lemurs to 1.5% isoflurane. Body temperature of mouse lemurs was continuously maintained at 36°–37°C using a heated pad connected to a controller that received feedback from a temperature sensor attached to the animal. Ag/AgCl gel-coated electrocardiogram electrodes (Unomedical; Herlev, Danimarca) were attached to the superior right and to the two inferior limbs of the mouse lemurs. The electrodes were connected to a standard one-lead electrocardiogram amplifier module (EMKA Technologies, Paris, France), which included high- and low-pass filters (set to 0.05 Hz and 500 Hz, respectively) and a gain selection device (set to 1,000-fold). Signals were digitized continuously at 2 kHz and recorded using an IOX data acquisition system (emka TECHNOLOGIES, Paris, France). The recordings were carried out for a 45-minute period, and the software ECGAuto (emka TECHNOLOGIES, Paris, France) was used to perform offline analysis of the recorded data. For each mouse lemur the mean HR, its standard deviation and the parameters characterizing the electrocardiogram wave, were calculated at 30-min intervals starting 10 min after the beginning of each 45-min recording.

### 4. Langendorff perfused hearts

After general anesthesia, consisting of 0.1 mg/g of ketamine (Imalgène, Merial, Bourgelat France) and complete loss of hind-limb reflex or any other body sensation, we removed the heart from the thoracic cage of mouse lemurs and quickly mounted it on a Langendorff apparatus (Isolated heart system; emka TECHNOLOGIES, Paris, France) at a constant pressure of 80 mm Hg with normal Tyrode’s solution. Perfused hearts were immersed in the water-jacked bath and maintained at 36°C. electrocardiograms were continuously recorded by Ag-AgCl electrodes positioned on the epicardial side of the right atrium close to the SAN and near the apex. Heart rate was allowed to stabilize for at least 30 min before perfusion of epinephrine (30 nM) or acetylcholine (50 nM).

### 5. Atrial preparation and optical voltage mapping

Atrial preparations (including the SAN and the right and left atria) were obtained as described previously.^46^ Briefly, after general anesthesia with ketamine and complete loss of hind-limb reflex (see above), we removed the heart from the thoracic cage of the animal. Then, we cut the coronary sinus and, starting from it, we removed the ventricles from the heart. We used a stereomicroscope (SZX16, Olympus; Tokyo, Japan) with low magnification (7X) to trans-illuminate and directly visualize the remaining atrial preparation. We identified the SAN region using the borders of the superior and inferior vena cava, the crista terminalis and the interatrial septum as landmarks.^33^ The atrial preparation was pinned to the bottom of an optical chamber (Fluorodish, FD35PDL-100, WPI; Sarasota, FL) coated with ~2 mm of clear Sylgard (Sylgard 184 Silicone elastomer kit; Dow Corning; Midland, MI). To avoid interference from secondary pacemaker tissues we removed the atrioventricular node from the preparation.

For comparative experiments, 24-month-old mice were used to match the old age of *Microcebus murinus* (from 6 to 11 years old) from which we obtain tissue preparation of the SAN. As for *Microcebus murinus* the investigation with mice conforms to the European directives (2010/63/EU) for the Care and Use of Laboratory Animals and was approved by the French Ministry of Agriculture (N° D34-172-13). Briefly, mice were anaesthetized with 0.01 mg/g xylazine (Rompun 2%, Bayer AG, Leverkusen Germany), 0.1 mg/g ketamine (Imalgène, Merial, Bourgelat France) and 0.2 mg/g Na-Pentobarbital (CEVA, France). Then, after complete loss of hind-limb reflex or any other sensation, we removed the heart from the thoracic cage of the animal and we further dissected it to obtain the entire atrial preparation including the SAN and the atria.^46^

To analyze voltage changes in the SAN preparation we loaded it by immersing the tissue in a Tyrode’s solution containing the voltage-sensitive indicator Di-4-ANEPPS (2 μmol/L; AAT Bioquest, Sunnyvale, California). This immersion was performed at room temperature (20-22°C) and lasted for at least 30 min. To maintain proper oxygenation, the chamber containing the tissue was maintained under agitation for the whole loading period. After the loading step, the tissue was washed in dye-free Tyrode’s solution for 15 min. During this step, we slowly increase the temperature to 34-36° C. The atrial preparation was then constantly superfused at 34-36° C and imaged by high-speed optical voltage mapping (1000 to 333 frames/s) on a MiCAM03 Camera - 256×256 pixel CMOS sensor, 17.6×17.6 mm (SciMedia; Costa Mesa, CA). This camera was mounted on a THT microscope, with two objectives (2X and 1.6X) that generated a field of view of 22 x 22 mm. A system constituted by a 150 W halogen light and a built-in shutter (SciMedia; Costa Mesa, California) was used as the excitation source of light for the voltage dye. The filter set included a 531/50 nm excitation filter, 580 nm dichroic mirror, and 580 long-pass emission filter. To avoid motion artefacts, we blocked mechanical activity using blebbistatin (10 μM; Tocris Bioscience; Bristol, UK).^46^ Optical raw data were analysed using dedicated software from the camera developer, BV workbench (Brainvision; Tokyo, Japan), in combination with ClampFit (ver. 10.0.7, Molecular Devices, LLC; San Jose, California).

### 6. Isolation of SAN myocytes

The SAN tissue of mouse lemur was isolated as described above and cut in tissue strips. Strips were then transferred into a low-Ca^2+^, low-Mg^2+^ solution containing (in mM): 140.0 NaCl, 5.4 KCl, 0.5 MgCl_2_, 0.2 CaCl_2_, 1.2 KH_2_PO_4_, 50.0 taurine, 5.5 d-glucose and 1.0 mg/ml BSA, plus 5.0 mM HEPES-NaOH (adjusted to pH 6.9 with NaOH). The tissue was enzymatically digested by adding 229 U/ml collagenase type II (Worthington Biochemical Corporation), 1.9 U/ml elastase (Boehringer Mannheim), 0.9 U/ml protease (Sigma-Aldrich), 1 mg/ml BSA, and 200 μM CaCl_2_. Tissue digestion was performed for a variable time of 9-13 min at 35°C with manual agitation using a flame-forged Pasteur pipette. Tissue strips were then washed and transferred into a medium containing (in mM): 70.0 L-glutamic acid, 20.0 KCl, 80.0 KOH, 10.0 (±) D-β-OH-butyric acid, 10.0 KH_2_PO_4_, 10.0 taurine, 1 mg/ml BSA, and 10.0 HEPES-KOH, adjusted at pH 7.4 with KOH. SAN myocytes were manually dissociated in KB solution at 35°C for about 10 min. Cellular automaticity was recovered by readapting the myocytes to physiological extracellular Na^+^ and Ca^2+^ concentrations by adding first a solution containing (in mM): 10.0 NaCl, 1.8 CaCl_2_, and subsequently normal Tyrode’s solution containing 1 mg/ml BSA. The final storage solution contained (in mM): 100.0 NaCl, 35.0 KCl, 1.3 CaCl_2_, 0.7 MgCl_2_, 14.0 l-glutamic acid, 2.0 (±)D-β -OH-butyric acid, 2.0 KH_2_PO_4_, 2.0 taurine, and 1.0 mg/ml BSA, pH 7.4. Cells were then stored at room temperature until use. All chemicals were obtained from Sigma-Aldrich, except for the (±)D-β - OHbutyric acid, which was purchased from Fluka Chemika. For electrophysiological recording, SAN myocytes in the storage solution were harvested in special custom-made recording Plexiglas chambers.

### 7. Preparing and culturing of MDSCs and PML cells

MDSCs were prepared using the modified preplate procedure originally described by Arsic et al.^37^ and modified recently in the reference.^38^ Briefly, in animals that had to be euthanized for natural aging or other factors hampering normal life, after general anesthesia, consisting of 0.1 mg/g of ketamine (Imalgène, Merial, Bourgelat France) and complete loss of hind-limb reflex or any other body sensation, the hind limbs were skinned and muscles were removed and stored in Hanks buffered saline overnight at 4°C. After mincing muscles were enzymatically dissociated at 37°C for 45 min in 0.2% collagenase A (Roche) and 1 mg/ml dispase (GibcoBRL). After digestion, muscle cells were passed through 100 μm and 40 μm nylon cell strainers (BD Falcon) and collected by centrifugation at 328 g for 10 min. Pelleted cells were washed twice in phosphate buffer saline (PBS), suspended in the growth medium [DMEM/F12 (Sigma), 16% fetal bovine serum (PAA), 1% Ultroser G (Pall Life Sciences), antibiotic-antimycotic mix (Gibco)] and plated on uncoated dishes (NUNC). Dishes were incubated overnight at 37°C and the next day non-adherent multipotent stem cells were transferred to a new dish, a process repeated every 24 h for 6 to 8 days. For confocal microscopy experiments, cells were seeded onto glass-bottom chambers (Fluorodish, FD35PDL-100, WPI; Sarasota, FL). Otherwise, they were cultured in standard 35 mm petri dishes. To ensure a full maturation of spontaneously differentiated stem cells, culturing was prolonged for 1 to 3 months before experimentation, and PML cells were selected as spontaneously-contractile cells, based on their morphology.

### 8. Patch-clamp recordings of *Microcebus murinus* SAN myocytes and PML cells

Myocytes isolated from the *Microcebus murinus* SANs were prepared as described in the previous sections, while PML cells were enzymatically treated with brief digestions, of 3 to 5 minutes, at 37 °C with 229 U/ml collagenase type II (Worthington Biochemical Corporation), and washed in Tyrode solution for enzyme removal, leaving cells in their culture petri-dish. The basal extracellular Tyrode’s solution used in all recordings contained (in mM): 140.0 NaCl, 5.4 KCl, 1.8 CaCl2, 1.0 MgC_l2_, 5.0 HEPES-NaOH, 5.5 and d-glucose (adjusted to pH 7.4 with NaOH). To measure the hyperpolarization-activated current *I_f_*, we employed standard Tyrode’s solution in the presence or absence of 2 mM Ba^2+^ to block the inward rectifier K^+^ current *I_k1_*, without affecting *I_f_*^60^. *I_k1_* was subtracted from *I_f_* as the net Ba^2+^-sensitive conductance, while *I_f_* was extracted as the currents not blocked under perfusion of Ba^2+^, but subsequently inhibited in the presence of 5 mM Cs^+^. *I_ca,L_* and *I_ca,T_* were recorded as previously described^49^. In particular, I_caToT_ (I_caT_ +I_caL_) was recorded from a holding potential of −80 mV in isolated SAN myocytes. Then after switching HP to −55 mV we inactivated I_caT_ to obtain only I_ca,L_.

AP firing activity was measured in the current clamp configuration. Patch-clamp electrodes had a resistance of 4-5 MΩ when filled with an intracellular solution containing (in mM): 130.0 K^+^-aspartate; 10.0 NaCl; 2.0 ATP-Na^+^ salt, 6.6 creatine phosphate, 0.1 GTP-Mg^2+^, 0.04 CaCl_2_ (pCa = 7.0), and 10.0 Hepes KOH (adjusted to pH 7.2 with KOH). For the modulation of AP firing, we perfused epinephrine (30 nM), acetylcholine (50 nM) or tetrodotoxin (30 μM). Compounds were added to the external medium and perfused through a pipette at a controlled temperature of 36°C. Pacemaker activity of SANs and PML cells were recorded under perforated patch conditions by adding β-escin (50 μM) to the pipette solution.

### 9. Measuring PML cell shortening

PML cells were placed on the stage of an inverted microscope (Olympus IX71). Cell contractions were recorded using a video-based edge-detection system (IonOptix™). Digitized images were collected at 120 Hz using an IonOptix™ Myocam-S CCD camera, and displayed within the IonWizard™ acquisition software (IonOptix™). To measure changes in cell length, two cursors were placed at opposite edges of the cell samples, where the difference in optical density (maximum: dark; minimum: light) generated the higher contrast. Cell shortening was then assessed by the simultaneous movement of the cursors. For the modulation of contraction rate, we perfused epinephrine (1 nM to 1 μM), acetylcholine (1 to 50 nM), ivabradine (3 nM to 3 μM) and E4031 (300 nM and 3 μM). Compounds were added to the external medium and perfused through a pipette at a controlled temperature of 36°C.

### 10. Immunofluorescence analysis

Immunofluorescence analysis was performed on SAN tissue, native SAN pacemaker myocytes and PML cells, as in the reference.^37^ Tissues were fixed in 4%paraformaldehyde for 6 hours while cells were fixed with 3.7% formaldehyde for 30 min. Then these samples were permeabilized for 30 s in - 20°C acetone, preincubated for 30 min with 0.5% BSA in PBS. Cells were then incubated for 1 h at 37°C with primary antibodies against HCN4, (1:100, Alomone), sarcomeric alpha-actinin, clone EA-53 (1:100, Sigma-Aldrich), connexin 45 (1:100, H-85, Santa Cruz Biotechnology), Ca_v_1.3 Ca^2+^ channel polyclonal antibody (N38.8, neuromab UC Davis), cardiac troponin I (1:100, Millipore), caveolin 3 (1:100, BD Transduction Laboratories) or myogenin (1:50, sc576 santa cruz). Specific staining was revealed using anti-mouse or anti-rabbit Alexa Fluor 488 and Alexa Fluor 555-conjugated secondary antibodies (InVitrogen) with Hoechst33358 nuclear stain (0,1 μg/ml) for 30 min at 37°C.

### 11. Statistical analysis

Significance was evaluated through paired or unpaired Student’s T test, one-way- and two-way ANOVA as specified in figure legends. When testing statistical differences, results were considered significant with p<0.05. Data analysis were performed with GraphPad Prism 9.0.

## Acknowledgments

We are thankful to the personnel of the animal facility of the lemur colony at Brunoy, to the animal facility at RAM-CECEMA, the MRI platform of imaging and the statistic platform Statabio. We thank Fu Lingjian, Dr. Yuquan Wang and Emma Zub for helping with software installation and exploitation. M-L DiFrancesco was recipient of a postdoctoral fellowship by the Laboratory of Excellence *Ion Channel Science and Therapeutics* supported by a grant from *Agence Nationale de la Recherche*(ANR-11-LABX-0015). This work was supported by ANR grants (2010-BLAN-1128-01 and ANR-15-CE14-0004-01 to M.E.M.) and the *Fondation Leducq* (TNE FANTASY 19CVD03 to Matteo E. Mangoni and Peter J. Mohler).

---

**Suppl. Figure 1:**
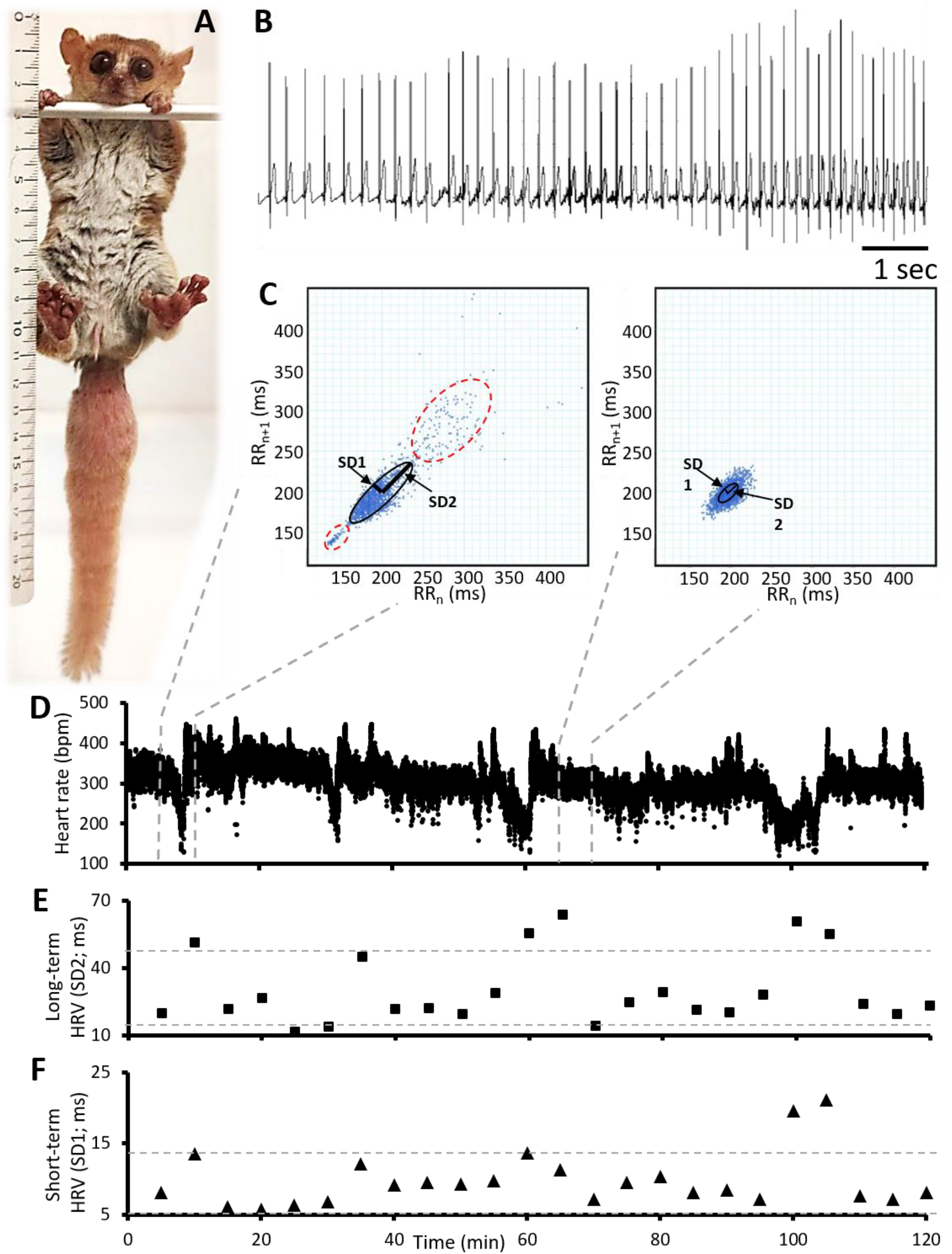
Indexes of long- and short-term heart rate variability in freely moving mouse lemurs. **A:** 4 years old female of grey mouse lemurs (*Microcebus murinus*). **B**: ECG recordings showing a very long RR interval followed by HR increase. **C:** Poincaré plots obtained analyzing intervals of 5 min during periods of high (left panel) and low (right panel) HRV in D. In both panels note the standard deviations from axis 1 and 2 (SD1 and SD2, respectively) which measure short- and long -term HRV, respectively. In the left panel note the presence of two clouds of points (red dotted ellipses) separated from the main one (black ellipse). The points belonging to the red dotted ellipse in the upper part of the panel correspond to slow HR (long RR intervals), while those belonging to the red dotted ellipse in the lower part of the panel correspond to fast HR (short RR intervals). Such subdivision is absent in the right panel, which represent by Poincaré an interval of 5 min with constant HR. **D:** HR versus time for a period of 120 min. **E** and **F:** Plots obtained by measuring the average long- and short-term HRV (SD2 and SD1, respectively) for intervals of 5 minutes of the recording in D. Dashed gray lines in E and F indicate the threshold of mean ± SD of the reported parameters in this panel.

**Suppl. Fig. 2:**
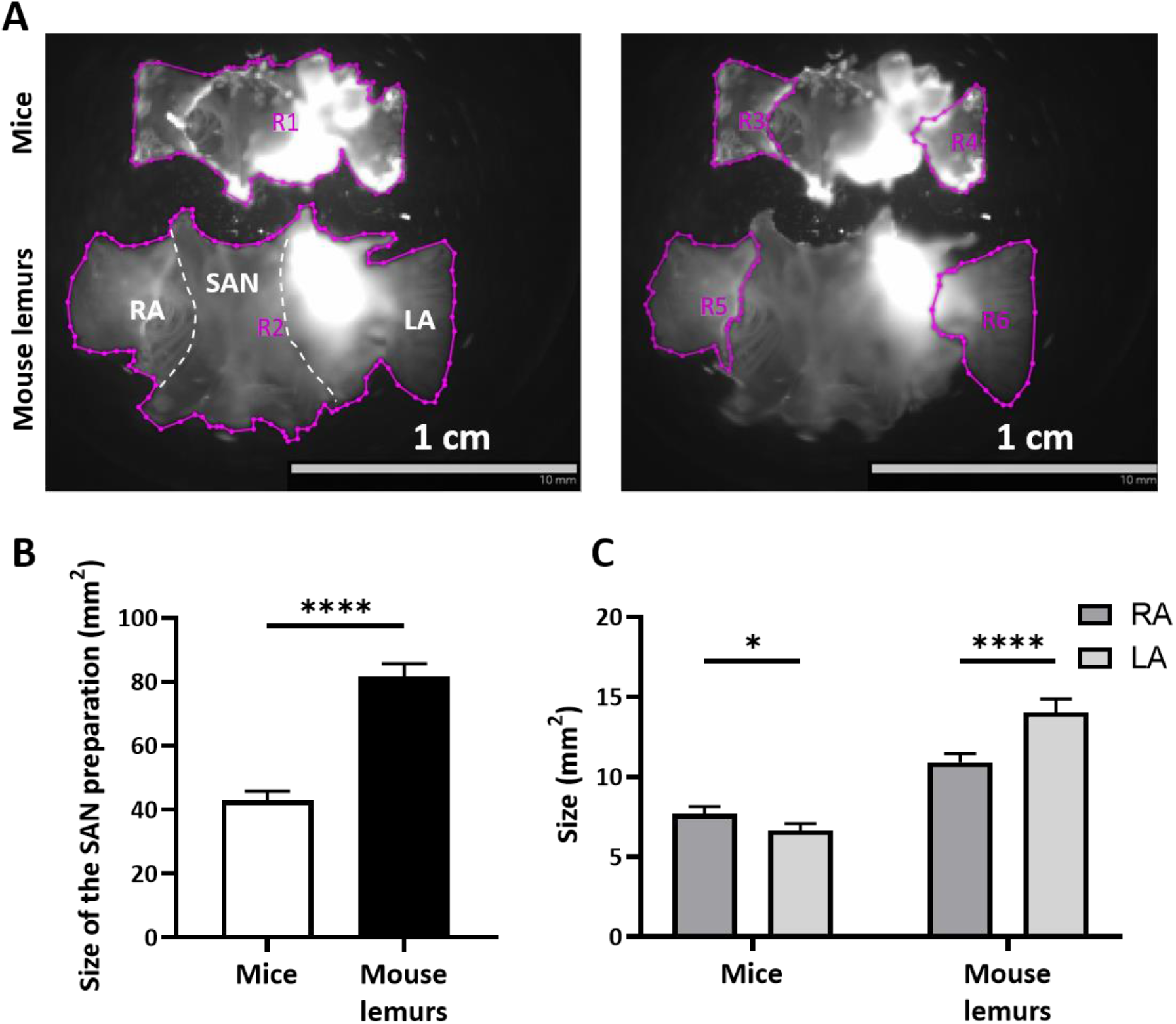
Comparative analysis of SAN-Atria preparations of mouse lemurs and mouse. **A:** SAN preparations of mice and mouse lemurs including SAN, right and left atria (RA and LA, respectively). Purple lines indicate the surface of the regions including the whole SAN preparation (R1 and R2 respectively), the surface of RA (R3 and R4, respectively) and the surface of LA in mice and mouse lemurs (R5 and R6, respectively). **B:** Size of the whole atrial preparations of mice and mouse lemurs represented in A. **C:** Size of RA and LA. N mice = 14 and n mouse lemurs = 11; *p<0.05, ****p<0.0001 by unpaired T-test and One-way Anova with Sidak’s multi-comparison test.

**Suppl. Video 1: Dynamics of pacemaker generation and conduction within the SAN preparation of mice (up) and mouse lemurs (down).** Green to red colors indicated the intensity of depolarization in arbitrary units of fluorescence.

